# Lethal giant larvae gene family (*Llgl1* and *Llgl2*) functions as a tumor suppressor in mouse skin epidermis

**DOI:** 10.1101/2023.03.06.531408

**Authors:** Victor M. Bii, Dmytro Rudoy, Olga Klezovitch, Valeri Vasioukhin

**Author notes:** Corresponding author: Valeri Vasioukhin (email).

## Abstract

Loss of cell polarity and tissue disorganization occurs in majority of epithelial cancers. Studies in simple model organisms identified molecular mechanisms responsible for the establishment and maintenance of cellular polarity, which play a pivotal role in establishing proper tissue architecture. The exact role of these cell polarity pathways in mammalian cancer is not completely understood. Here we analyzed the mammalian orthologs of drosophila apical-basal polarity gene lethal giant larvae (*lgl*), which regulates asymmetric stem cell division and functions as a tumor suppressor in flies. There are two mammalian orthologs of *lgl* (*Llgl1* and *Llgl2*). To determine the role of the entire lgl signaling pathway in mammals we generated mice with ablation of both *Llgl1* and *Llgl2* in skin epidermis using K14-Cre (*Llgl1/2*^*-/-*^ cKO mice). Surprisingly, we found that ablation of *Llgl1/2* genes does not impact epidermal polarity in adult mice. However, old *Llgl1/2* cKO mice present with focal skin lesions which are missing epidermal layer and ripe with inflammation. To determine the role of lgl signaling pathway in cancer we generated *Trp53*^*-/-*^*/Llgl1/2*^*-/-*^ cKO and *Trp53*^*-/+*^*/Llgl1/2*^*-/-*^ cKO mice. Loss of *Llgl1/2* promoted squamous cell carcinoma (SCC) development in *Trp53*^*-/-*^ cKO and caused SCC in *Trp53*^*-/+*^ cKO mice, while no cancer was observed in *Trp53*^*-/+*^ cKO controls. Mechanistically, we show that ablation of *Llgl1/2* causes activation of aPKC and upregulation of NF-kB signaling pathway, which may be necessary for SCC in *Trp53*^*-/+*^*/Llgl1/2*^*-/-*^ cKO mice. We conclude that Lgl signaling pathway functions as a tumor suppressor in mammalian skin epidermis.

## INTRODUCTION

Due to the extraordinary progress in cancer research, significant information was acquired about the genetic modifications in human cancer. However, the early cellular events that trigger cancer are not well understood. It is likely that the defective asymmetric cell divisions of stem or progenitor cells are ultimately responsible for the accumulation of dividing cells that do not withdraw from the cell cycle and form a tumor (Bajaj et al. 2015). The process of normal asymmetric cell division of stem cells is responsible for formation of two daughter cells with different cell fates (Horvitz and Herskowitz 1992; Knoblich 2010). One cell continues to be a stem cell while the other cell undergoes differentiation and eventually withdraws from the cell cycle. It is plausible that the failure of proper asymmetric cell division transforms normal stem cell into a cancer stem cell that can cause tumor formation. To understand such earlier cancer initiation events, it is necessary to understand the mechanisms regulating asymmetric cell divisions and what happens to these mechanisms in cells that acquired genetic alterations that ultimately result in cancer development.

Most of the knowledge about the mechanisms of asymmetric cell division comes from studies on simple model organisms. Drosophila gene *lethal (2) giant larvae* (*lgl*) plays a critical role in asymmetric cell division of neuroblasts (Vasioukhin 2006). *L(2)gl* mutant flies show a dramatic cancer-like phenotype (Bilder et al. 2000). The brain and imaginal discs of *l(2)gl* larvae expand and the giant larvae dies (thus the name – lethal giant larvae) (De Lorenzo et al. 1999). Lgl functions in asymmetric cell division as a cell polarity protein, which is necessary for polarization of neuroblasts before cell division. Lgl regulates cell polarity by promoting the identity of the lateral and basal membrane domains and inhibiting the function of the Par6-Par3 (Bazooka)-aPKC and Crumbs-Stardust-Patj protein complexes responsible for the maintenance of the apical membrane domain (Bilder et al. 2000; Bilder et al. 2003; Tanentzapf and Tepass 2003). During neuroblasts division, apically localized Par3/6-aPKC-Pins-Ga/bg-Inscutable proteins orient the mitotic spindle and segregate the cell fate determinants Numb, Prospero and Brat to the basal cortex. After the division, cell fate determinants are present in only one daughter cell, which starts the differentiation program. Lgl is the principal downstream phosphorylation target of apical aPKC and mitotic Aurora A protein kinases (Betschinger et al. 2003; Rolls et al. 2003; Bell et al. 2015; Carvalho et al. 2015). aPKC and Aurora A phosphorylate Lgl and this phosphorylation releases Lgl from its association with membranes and actin cytoskeleton (Dong et al. 2015). In *lgl* mutant neuroblasts the cell fate determinants are not localizing basally and they are inherited by both daughter cells (Ohshiro et al. 2000; Peng et al. 2000). *Lgl* mutant neuroblasts and their progeny lose their ability to differentiate and continue to proliferate (Woods and Bryant 1989). This results in massive hyperplasia in the brain of zygotic *lgl* mutant flies (Gateff and Schneiderman 1969).

The mechanisms responsible for *lgl* function in Drosophila are complex and not completely understood. In Drosophila neuroblasts, loss of *lgl* results in increased activity of aPKC, which is necessary and sufficient for hyperplasia (Rolls et al. 2003; Lee et al. 2006). There is significant evidence that *lgl* is necessary for the regulation of the tumor-suppressive Hippo pathway and the Hippo effector Yorkie (Yap1 ortholog) is hyperactive in *lgl*-mutant flies (Grzeschik et al. 2010; Parsons et al. 2010; Enomoto and Igaki 2011; Jukam and Desplan 2011; Parsons et al. 2014a). In addition, Drosophila *lgl* negatively regulates Notch and JNK signaling (Sun and Irvine 2011; Parsons et al. 2014b). It is necessary for the appropriate cargo sorting into the retromer pathway (de Vreede et al. 2014), and endosomal vesicle acidification (Portela et al. 2018). Recent RNAi kinome screen in Drosophila not only confirmed the connection between lgl and aPKC, Hippo, Notch and JNK, but also identified genetic interactions between lgl and Dpp, Wg, Hh, Src, Ras, and PI3K signaling pathways, indicating that lgl-mediated tumor suppressive mechanisms in Drosophila can be very complex and cell context dependent (Parsons et al. 2017).

There are two mammalian orthologs of Drosophila *lgl* gene, *Llgl1* and *Llgl2*, which are expressed in largely overlapping pattern (Klezovitch et al. 2004). The expression of LLGL1/2 proteins is lost in variety of human cancers including gastric cancer (Nam et al. 2014), lung squamous cell carcinoma (Matsuzaki et al. 2015), hepatocellular carcinoma (Lu et al. 2009), glioma (Liu et al. 2015), endometrial cancer (Tsuruga et al. 2007), and melanoma (Kuphal et al. 2006). Re-expression of LLGL proteins in cancer cell lines that lost endogenous LLGL1/2 results in decreased cell proliferation and survival and increased differentiation (Schimanski et al. 2005; Kuphal et al. 2006; Kashyap et al. 2013; Gont et al. 2014; Greenwood et al. 2016). Interestingly, in human glioblastoma cells LLGL1 is a critical regulator of self-renewal and differentiation that functions downstream from PTEN (Gont et al. 2013). In these cells, absence of PTEN promotes self-renewal and inhibits differentiation, and the mechanisms involve activation of aPKC and inhibition of LLGL1 by aPKC-mediated phosphorylation (Gont et al. 2013; Gont et al. 2014). Expression of mutant LLGL1 (LLGL1-SA) that cannot be phosphorylated by aPKC completely erases the PTEN-loss-mediated self-renewal and differentiation phenotypes (Gont et al. 2013; Gont et al. 2014). In oligodendrocyte progenitors, *Llgl1* regulates differentiation and cooperates with *Ink4a/Arf* in transformation and gliomagenesis (Daynac et al. 2018). We have previously used mouse genetic approach to determine the role of *Llgl1 and Llgl2* genes in mammalian organism in vivo. We found that mice with null mutation of *Llgl2* display attenuated placental branching morphogenesis but show no obvious adult phenotypes (Sripathy et al. 2011). In contrast, mice with germline ablation of *Llgl1* display prominent disorganization of the developing brain and die within few hours after birth from severe hydrocephalus (Klezovitch et al. 2004). Conditional deletion of *Llgl1* in embryonic brain revealed a connection between Llgl1 and accumulation of N-cadherin at the apical junctional complexes (Jossin et al. 2017). The loss of Llgl1 resulted in disruption of epithelial adhesion and brain malformation; however, no cancer was observed in these animals (Jossin et al. 2017). Since *Llgl1* and *Llgl2* are very similar to each other, we hypothesized that these genes may display genetic redundancy in mice organisms and the ablation of both genes in the same tissue is necessary to reveal the role of the LGL pathway in mammals. We have now generated the conditional double mutant *Llgl1/2* mice and this approach revealed tumor suppressive role of *Llgl1/2* in mammalian skin epidermis.

## RESULTS

### Generation of epidermis–specific double mutant *Llgl1/Llgl2* conditional knockout (cKO) mice (*K14-Cre/Llgl1/2*^*fl/fl*^)

We have previously generated mice with a null mutation of *Llgl1* and found that this mutation results in neonatal lethal phenotype (Klezovitch et al. 2004). *Llgl1*^*-/-*^ pups develop disorganization of brain architecture; however, the phenotype is confined to brain because it is the only embryonic tissue that does not express *Llgl2* (Klezovitch et al. 2004). Our work on *Llgl2* mutant mice revealed viability and fertility of straight *Llgl2* knockout mice, presumably due to compensatory expression of *Llgl1* in all tissues (Sripathy et al. 2011). We reasoned that to understand the role of the entire Lgl signaling pathway in cancer we need to generate mice with Cre-inducible tissue-specific conditional double knockout of *Llgl1* and *Llgl2*. Our previously published *Llgl2* knockout mice were made using *Llgl2* allele gene trapped with vector pGT0lxf (Sanger Institute), which reverts to wild-type phenotype upon Cre recombination. Thus, we started this work by engineering mice with new *Llgl2* mutation which generates strong loss-of-function allele upon Cre-mediated recombination (Supplemental Fig.1). We then crossed these mice with our previously generated *Llgl1*^*flox/flox*^ mice (Jossin et al. 2017). Both *Llgl1* and *Llgl2* genes reside on mouse chromosome 11. Therefore, we relied on genetic crossing over to obtain mice with double targeted *Llgl1* and *Llgl2* alleles. To determine the role of the entire Lgl pathway in adult mouse tissue homeostasis and tumor suppression, we decided to make mice with deletion of both *Llgl1/2* genes in skin epidermis utilizing our previously generated and characterized Keratin14-Cre mice (Vasioukhin et al. 1999).

Unexpectedly, we found that inactivation of both *Llgl1* and *Llgl2* genes in skin epidermis gave rise to mice that were live and fertile. Western blot analyses and immunofluorescent stainings revealed loss of both LLGL1 and LLGL2 proteins in skin epidermis of *Llgl1/2* cKO mice (Fig. 1A-C). We did not observe tumor development in these animals; however, we noticed that many older *Llgl1/2* cKO mice lose hair and develop prominent wound–like lesions riddled with inflammation (Fig. 1D-F’). Deletion of many tumor-suppressor genes does not cause frank tumor development and their cancer-relevant functions becomes apparent only when these mutations are analyzed in tissues already predisposed to cancer development. Therefore, we decided to analyze *Llgl1/2* in cancer sensitized genetic background using conditional deletion of one allele of *Trp53*.

**Fig. 1.**
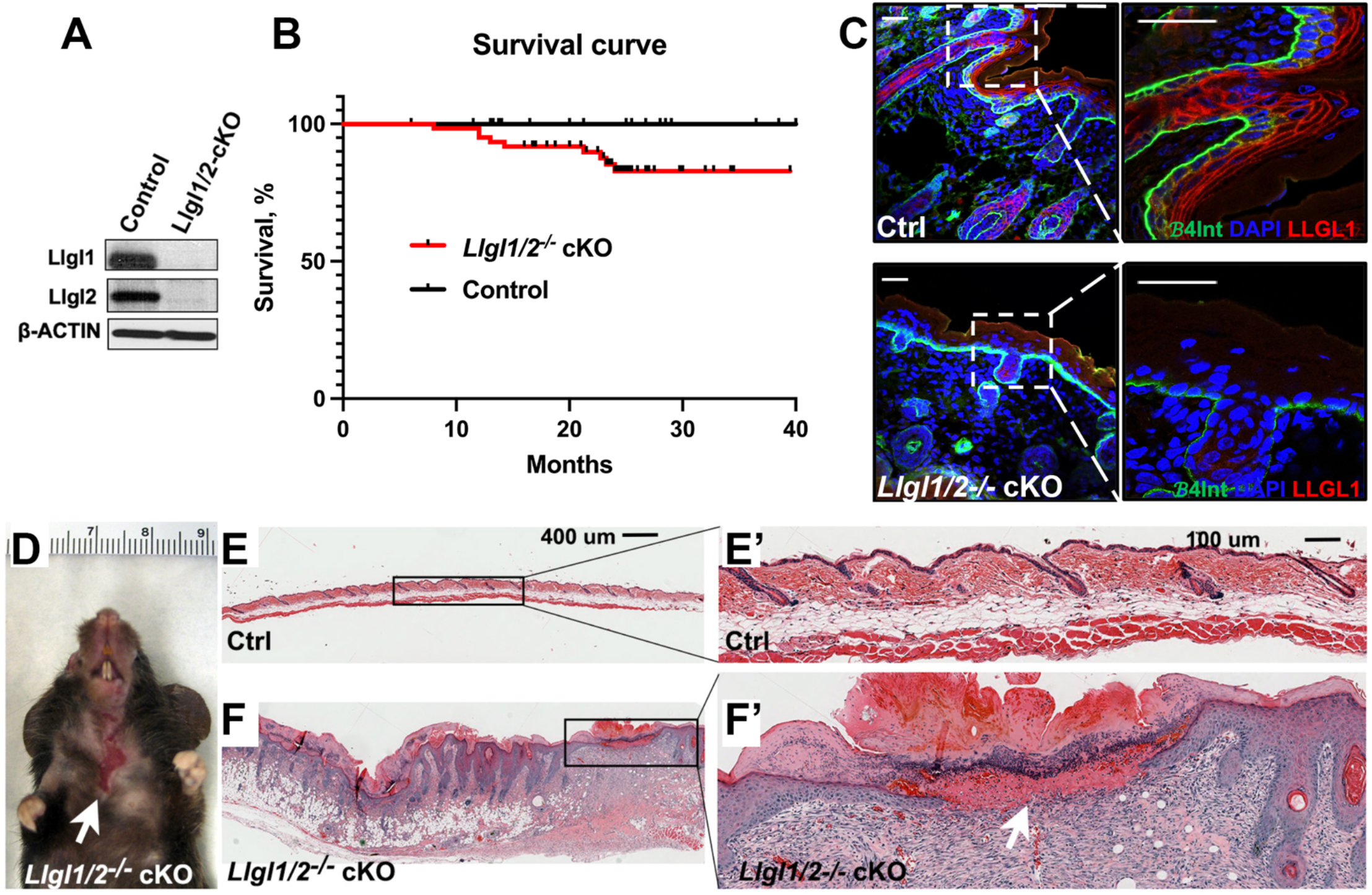
Generation of mice with epidermis-specific knockout of *Llgl1* and *Llgl2* (*Llgl1/2*^*-/-*^ cKO mice). **A**. Western blot analysis of proteins from epidermises of newborn mice with indicated genotypes using anti-Llgl1 and anti-Llgl2 antibodies. **B**. Survival curve of Control (K14-Cre) and *Llgl1/2*^*-/-*^ cKO (K14-Cre/*Llgl1/2*^*fl/fl*^) mice. Log-Rank test p=0.07. **C**. Confocal images of immunofluorescent staining of skin sections from Control (K14-Cre) and *Llgl1/2*^*-/-*^ cKO (K14-Cre/*Llgl1/2*^*fl/fl*^) new-born pups with anti-β4 integrin (green) and anti-LLGL1/2 (red) antibodies. Areas in black boxes are shown at higher magnification on the right. DAPI (blue) indicates nuclear counter stain. Scale bar 10μm. **D**. Loss of hair and prominent wound like lesions in 1 year old *Llgl1/2*^*-/-*^ cKO individual. **E-F’**. H&E staining of skin sections from K14-Cre (Ctrl) and K14-Cre/*Llgl1/2*^*fl/fl*^ (*Llgl1/2*^*-/-*^ cKO) mice. Images in boxes in E and F are shown at higher magnification in E’ and F’. Same magnification for E-F and E’-F’. Note prominent inflammation and lesion missing the epidermal skin layer in 1 year old *Llgl1/2*^*-/-*^ cKO individual (white arrows in D and F’).

### Loss of *Llgl1/2* genes in *Trp53*^*+/-*^ cancer sensitized genetic background (*K14-Cre/Llgl1/2*^*fl/fl*^*/Tp53*^*fl/+*^ mice) results in highly penetrant development of skin SCCs, while cancer is extremely rare in *K14-Cre/ Tp53*^*fl/+*^ mice

*Trp53* is the most frequently inactivated tumor-suppressor in mammalian genome and it is frequently mutant in skin tumors. Mouse *Trp53* resides on the same chromosome as *Llgl1/2*, and we again used crossing over to obtain mice with triple conditional *Llgl1/2/Trp53* genes. These mice were used to generate double knockout of *Llgl1 and Llgl2* and knockout of single allele of *Trp53* (*K14-Cre/Llgl1/2*^*fl/fl*^*/ Tp53*^*fl/+*^ cKO mice). *K14-Cre/Tp53*^*fl/+*^ cKO animals were used as controls. We found that while mice with deletion of *Llgl1/2* and one allele of *Trp53* develop skin SCC tumors with a half-time of ∽13 months, no tumor development was observed in control *K14-Cre/Tp53*^*fl/+*^ cKO animals (Fig. 2). Mice with homozygous deletion of *Trp53* (*K14-Cre/Tp53*^*fl/fl*^ mice) develop SCC tumors; however, simultaneous deletion of *Llgl1/2* results in much earlier development of SCC tumors in *K14-Cre/ Llgl1/2*^*fl/fl*^*/Tp53*^*fl/fl*^ mice (p-value<0.0001,Supplementary Fig.2). These data provide strong genetic proof of tumor suppression function of *Llgl1/2*-mediated signaling pathway in mouse epidermis. Moreover, since we observed a binary (tumors present, yes/no) phenotypic difference between *K14-Cre/Llgl1/2*^*fl/fl*^*/ Tp53*^*fl/+*^ cKO and control *K14-Cre/ Tp53*^*fl/+*^ cKO animals, these mutant mice combination provided us with an excellent in vivo model system to dissect the role of Lgl pathway in tumor suppression.

**Fig. 2.**
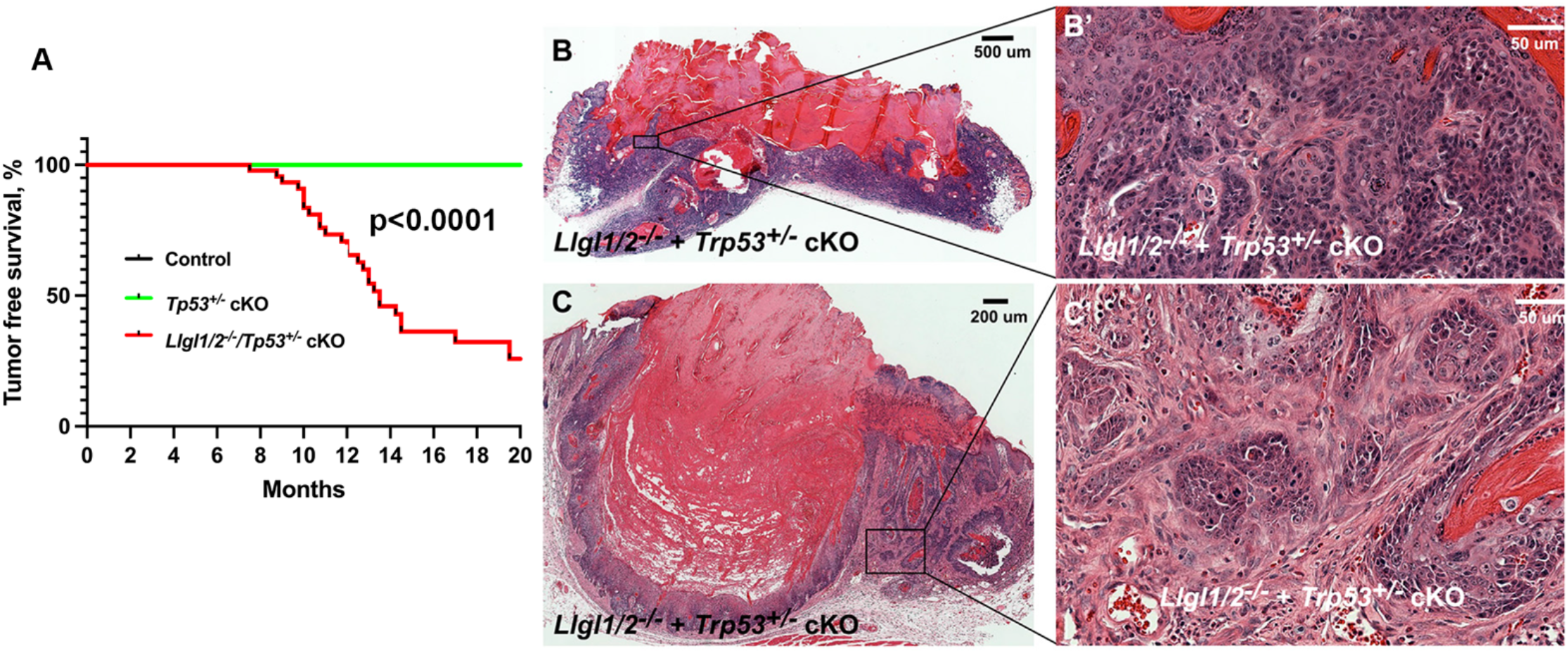
Tumor suppressor function of *Llgl1/2* in cancer sensitized heterozygous *Trp53*^*+/-*^ genetic background. **A**. Survival curve of Control (K14-Cre), *Trp53*^*+/-*^ cKO (*K14-Cre/Trp53*^*+/fl*^) and *Llgl1/2*^*-/-*^*/Trp53*^*+/-*^ cKO (*K14-Cre/Llgl1/2*^*fl/fl*^*/Trp53*^*+/fl*^) mice. Log-Rank test. **B-C’**. H&E staining of SCC skin tumors from *K14-Cre/Llgl1/2*^*fl/fl*^*/Trp53*^*+/fl*^ mice. Images in boxes in B and C are shown at higher magnification in B’ and C’.

### Generation of *Llgl1/2/Trp53* mutant and control primary mouse keratinocyte cell lines

To analyze the mechanisms of Lgl tumor-suppressive pathway in mouse epidermis, we decided to initiate the biochemical studies using cultured primary keratinocytes. Traditionally in the field, long-term cultures of primary mouse keratinocytes are established in low Ca^2+^ E-media in the presence of NIH3T3 feeder cells (Vasioukhin et al. 2000). However, low Ca^2+^ conditions are highly artificial, as keratinocytes cannot form cell-cell junction structures and develop cohesive epithelial cell layer, which is present in skin epidermis in vivo. Therefore, we investigated various primary keratinocyte culture conditions and found that normal Ca^2+^ organoid cell culture media developed in Dr. Hans Clevers’ laboratory (Karthaus et al. 2014) combined with coating of tissue culture plastic surface with laminin (Sigma) supports the growth, passaging and extensive expansion of primary mouse keratinocytes (Fig. 3). We used these conditions to establish primary cultures of keratinocytes from *Llgl1/2* ^*fl/fl*^, *Llgl1/2* ^*fl/fl*^ */Tp53*^*fl/+*^ *and Trp53*^*fl/+*^ mice. Infection of these cells with either Adenovirus-Cre-GFP or control Adenovirus-GFP particles generates genetic-background-matched pairs of mutant and control cell populations (Fig. 3A-C).

**Fig. 3.**
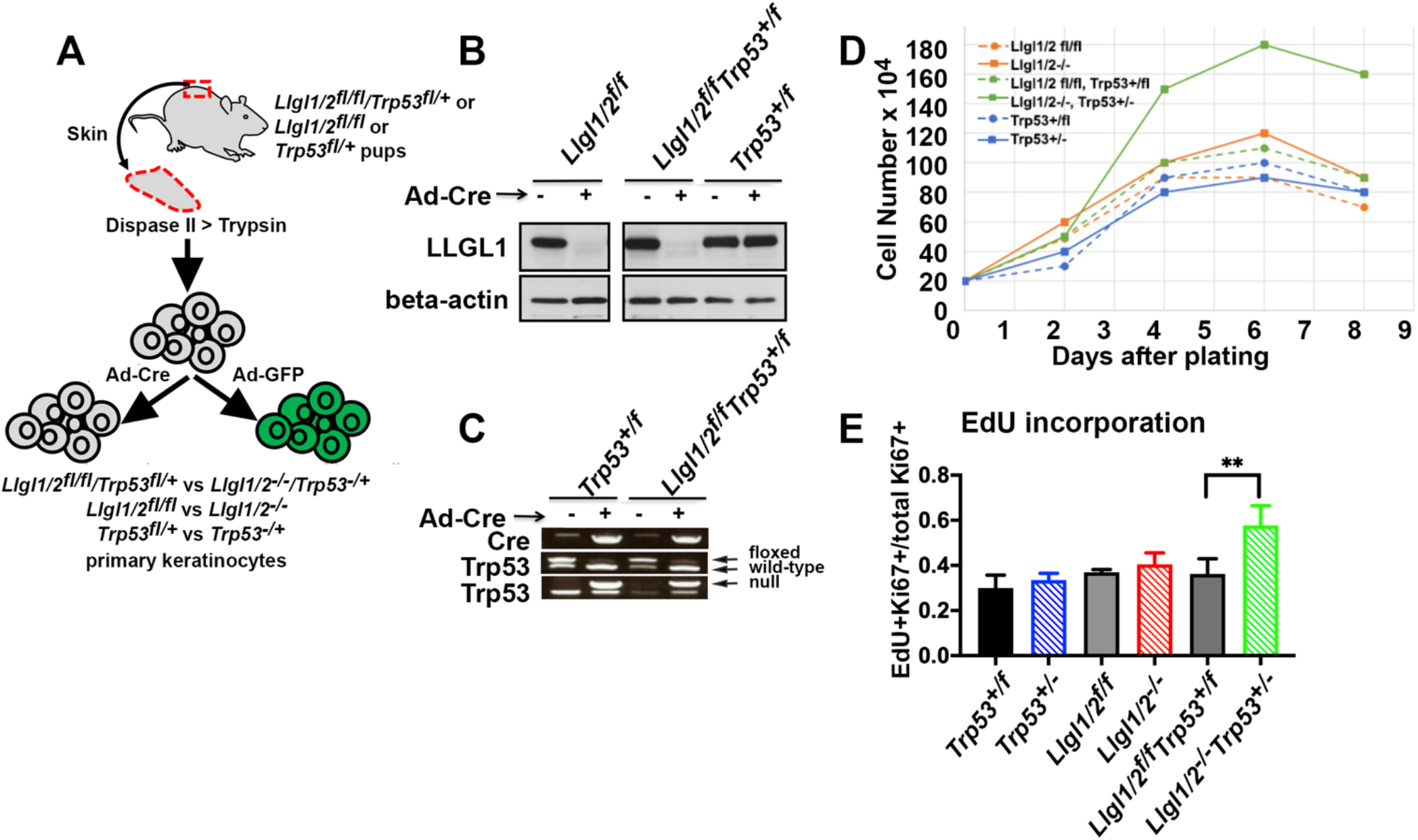
Loss of *Llgl1/2* cooperates with deletion of one allele of *Trp53* to promote proliferation and colony formation ability in primary keratinocytes. **A**. Generation of indicated mutant and matched control primary keratinocytes using tissue isolation and transient infections with Ad-GFP (control) and Ad-Cre-GFP adenoviruses. **B**. Western blot analysis of primary keratinocytes with indicated genotypes using anti-LLGL1/2 and anti-beta-actin antibodies. **C**. PCR analysis of primary keratinocytes with indicated genotypes using oligos amplifying *Cre, Trp53* wild-type, floxed and mutant alleles. **D**. Growth curves of primary keratinocytes with indicated genotypes. Graph shows mean values from 4 separate wells. **E**. Quantitation of one hour pulse EdU incorporation of primary keratinocytes with indicated genotypes. Graph shows mean values with standard deviation. ** -indicates p value <0.01 (Student’s t-test).

### Primary *Llgl12*^*-/-*^*/Tp53*^*+/-*^ keratinocytes display prominent increase in cellular accumulation

*To* determine whether *Llgl1/2*^-/-^*/Trp53*^-/+^ cells display any phenotypes relevant to cellular transformation that can help to determine the mechanism of *Llgl1/2* in tumor suppression, we analyzed *Llgl1/2*^-/-^, *Llgl1/2*^-/-^*/Tp53*^-/+^ *and Trp53*^-/+^ keratinocytes and their corresponding controls. We performed growth curve analyses and found that *Llgl1/2*^-/-^*/Tp53*^-/+^, but not their corresponding control, *Llgl1/2*^-/-^ or *Trp53*^-/+^ cells, display prominent increase in cellular accumulation (Fig. 3D). Similarly, *Llgl1/2*^-/-^*/Tp53*^-/+^ cells showed prominent increase in EdU incorporation (Fig. 3E). These tumor-relevant phenotypes resemble the situation in vivo, where we see tumor development in *Llgl1/2*^-/-^*/Trp53*^-/+^, but not in *Llgl1/2*^-/-^ or *Trp53*^-/+^ cKO mice (Fig. 2). Therefore, we decided to utilize primary cultures of *Llgl1/2*^-/-^*/Tp53*^-/+^ keratinocytes and their corresponding controls to reveal the mechanisms of *Llgl1/2* mediated tumor suppression.

### Activation of NF-kB signaling pathway in *Llgl1/2*^*-/-*^ primary keratinocytes in culture

To obtain initial insights into the *Llgl1/2* function in tumor suppression, we analyzed cell-autonomous transcriptional changes in *Llgl1/2*^-/-^ primary keratinocytes. RNA-seq experiments revealed substantial changes in gene expression in *Llgl1/2*^*-/-*^ cells (Fig. 4). Unbiased Gene Set Enrichment Analysis (GSEA) of these changes using “Hallmark Gene Sets” revealed activation of interferon alpha and beta signaling as two most significantly enriched gene sets (Fig. 4A). This was surprising because LLGLs have not been previously associated with innate immune signaling pathways. To continue our analysis of signaling pathways in *Llgl1/2*^-/-^ cells, we performed Western blot experiments with proteins extracted from cultured primary keratinocytes. Since Lgl has been previously strongly implicated in regulation of Hippo signaling pathway (Jukam and Desplan 2011; Parsons et al. 2014a; Li et al. 2017), which plays a very important tumor suppressive role in skin epidermis (Silvis et al. 2011; Li et al. 2016), we analyzed potential changes in YAP1 levels and its inhibitory S127 phosphorylation (Fig. 4B). Surprisingly, we did not observe a decrease but rather a mild increase in S127 specific phosphorylation of YAP1 (P-S127 YAP1/total YAP1) in *Llgl1/2*^-/-^ primary keratinocytes (Fig.4B). Since our RNA-Seq experiments detected cell autonomous changes in interferon signaling in *Llgl1/2*^-/-^ keratinocytes (Fig. 4A), we analyzed NF-kB signaling pathway. Consistent with data from transcriptional analysis, Western blot experiments revealed prominent increase in specific phosphorylation of Ser536 of RelA (NF-kB p65) (Fig. 4B). Specific phosphorylation of IKK-beta, an NF-kB kinase, which phosphorylates RelA at S536, was also increased in *Llgl1/2*^-/-^ keratinocytes (Fig. 4B). These data revealed activation of NF-kB signaling in *Llgl1/2*^-/-^ keratinocytes. The NF-kB signaling activation was specific to cells with deletion of Llgl1/2, as it was observed in both *Llgl1/2*^-/-^and *Llgl1/2*^-/-^*/Trp53*^-/+^ but not in *Tp53*^-/+^ keratinocytes (Fig. 4B).

**Fig. 4.**
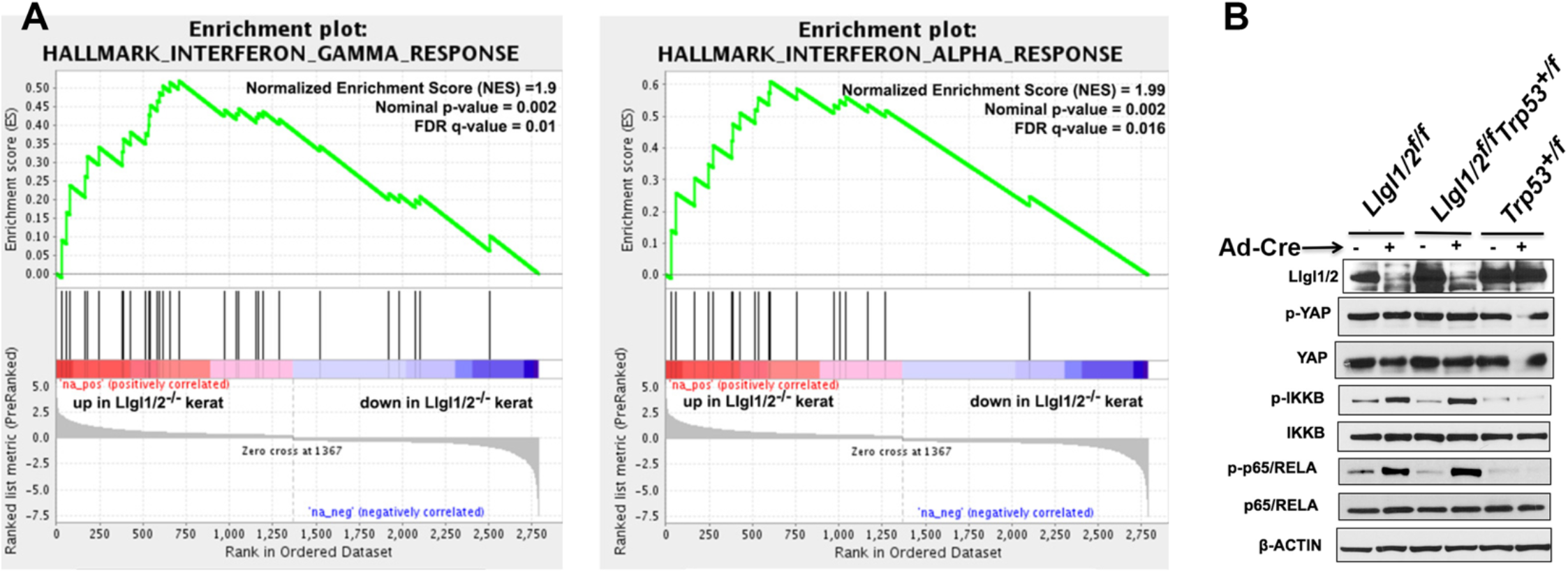
Activation of NFkB signaling pathway in *Llgl1/2*^*-/-*^ cells. **A**. RNAs extracted from *Llgl1/2*^*fl/fl*^ and *Llgl1/2*^*-/-*^ primary keratinocytes was analyzed by RNA-Seq (3 samples for each cell type). Significant differences in gene expression (q<0.05) were investigated by Gene Set Enrichment Analysis (GSEA). Two most significantly enriched Hallmark gene sets in *Llgl1/2*^*-/-*^ keratinocytes are Interferons gamma and alpha response sets (q=0.01 and q=0.016). **B**. Western blot analysis of proteins extracted from primary keratinocytes with indicated genotypes. Note increase in phosphorylation of IKKbeta and RelA in *Llgl1/2*^*-/-*^ and *Llgl1/2*^*-/-*^*/Trp53*^*+/-*^, but not in *Trp53*^*+/-*^ primary keratinocytes.

### Activation of NF-kB signaling pathway in *Llgl1/2*^*-/-*^ epidermis in vivo

Since we found activation of NFkB signaling pathway in *Llgl1/2*^*-/-*^ keratinocytes in culture, we thought to determine whether this signaling pathway is also activated in *Llgl1/2*^*-/-*^ cKO skin epidermis in vivo. For this purpose we isolated epidermises from *K14-Cre/Llgl1/2*^*fl/fl*^, *K14-Cre/Llgl1/2*^*fl/fl*^*/Tp53*^*fl/+*^ and control *K14-Cre/ Tp53*^*fl/+*^ cKO mice and analyzed them using RNA-Seq and Western blotting techniques. Similar to our findings in cultured keratinocytes, RNA-Seq and Western blot experiments revealed increase in activating phosphorylations of RelA (P-Ser536) and IKKbeta (P-Ser180) indicating activation of NFkB signaling pathway in epidermises from *Llgl1/2*^*-/-*^ *and Llgl1/2*^*-/-*^*/Trp53*^*-/+*^ but not from control *Trp53*^*-/+*^ cKO mice (Fig.5). We also found an increase in activating phosphorylation of atypical PKCiota (aPKCi) in *Llgl1/2*^*-/-*^ epidermis (Fig. 5B). This is consistent with previous findings reporting strong interaction between LLGL proteins and aPKC and negative regulation of aPKC by LLGLs (Almagor et al. 2019; Scott et al. 2019). aPKC can directly phosphorylate and activate IKK-beta at Ser180 and activated IKK-beta directly phosphorylates and activates RelA at Ser536 (Lallena et al. 1999). Thus, LLGL1/2-loss mediated activation of aPKC can result in activation of IKKbeta and the entire NFkB signaling pathway.

**Fig. 5.**
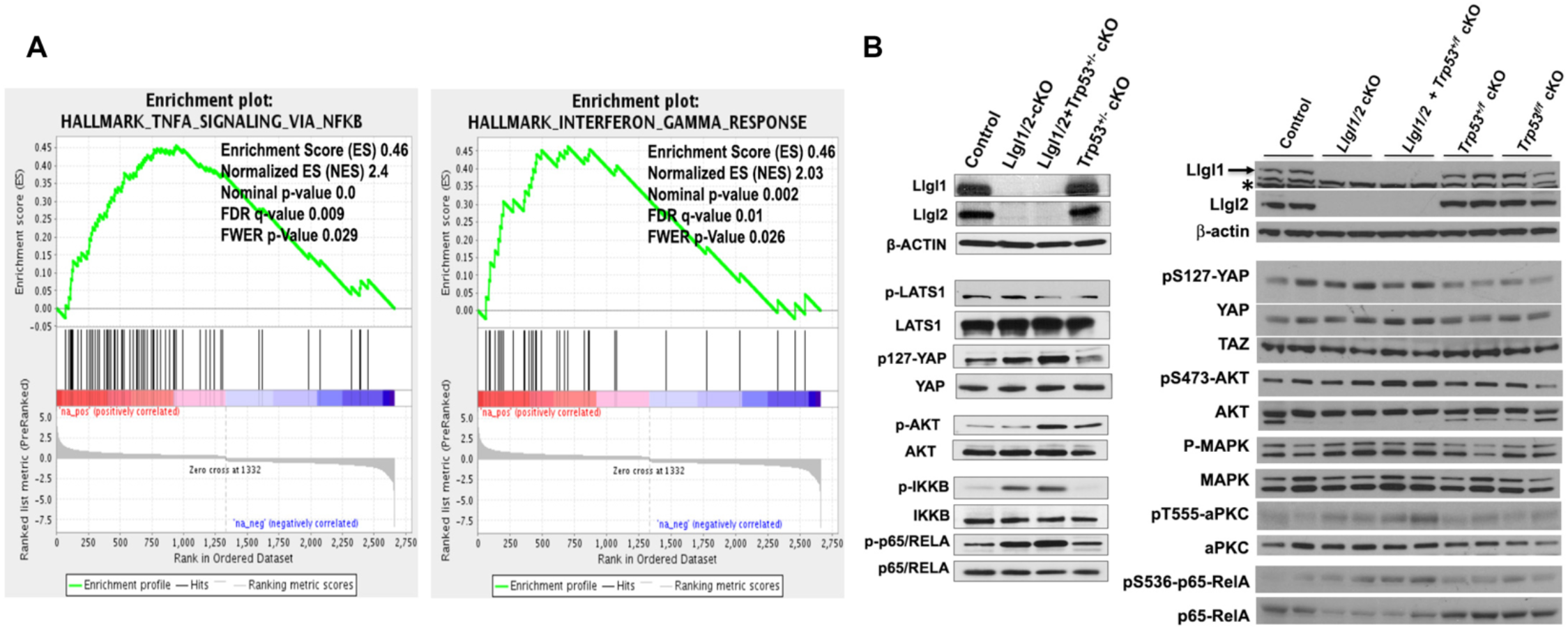
Activation of NFkB signaling pathway in *Llgl1/2*^*-/-*^ cKO epidermis. **A**. RNAs extracted from p0 Control (*K14-Cre*) and *Llgl1/2* cKO (K14-Cre/*Llgl1/2*^*fl/fl*^) skin epidermises was analyzed by RNA-Seq (3 individuals for each genotype). Significant differences in gene expression (q<0.05) were investigated by Gene Set Enrichment Analysis (GSEA). Significantly enriched Hallmark gene sets in *Llgl1/2-/-*epidermises are Interferon gamma and TNFA signaling via NFkB sets (q=0.01 and q=0.009). **B**. Western blot analyses of proteins extracted from epidermises of newborn mice with indicated genotypes. Note increase in phosphorylation of IKKbeta, RelA and aPKC in *Llgl1/2*^*-/-*^ cKO and *Llgl1/2*^*-/-*^ */Trp53*^*+/-*^ cKO, but not in *Trp53*^*+/-*^ cKO epidermises. * Indicates non-specific background protein band.

### RelA and aPKCi are necessary for hyperproliferation of *Llgl1/2*^-/-^*/Trp53*^-/+^ keratinocytes

We hypothesized that increased aPKC-NFkB activity can cooperate with the loss of one allele of *Trp53* to induce SCC tumor formation in *Llgl1/2*^*-/-*^*/Trp53*^*-/+*^*cKO* epidermis. To begin to address this hypothesis we analyzed the significance of these signaling pathways in hyperproliferation phenotype of *Llgl1/2*^-/-^*/Trp53*^-/+^ keratinocytes. RelA is a critical and one of the most abundant NF-kB subunits in skin epidermis (Poligone et al. 2013). To determine the significance of RELA we inactivated *RelA* in *Llgl1/2*^*-/-*^*/Trp53*^*-/+*^ and control *Llgl1/2*^*fl/fl*^*/Tp53*^*fl/+*^ keratinocytes. Knockdown of RelA with two independent shRNA constructs erased the differences in EdU incorporation between *Llgl1/2*^*-/-*^*/Trp53*^*-/+*^ and control *Llgl1/2*^*fl/fl*^*/Tp53*^*fl/+*^ cells (Fig. 6A,B). Mutation of *RelA* using two independent CRISPR/Cas9 constructs resulted in similar phenotype (Fig. 6C, D). We conclude that RelA activity is necessary for hyperproliferation phenotype of *Llgl1/2*^-/-^*/Trp53*^-/+^ keratinocytes.

**Fig. 6.**
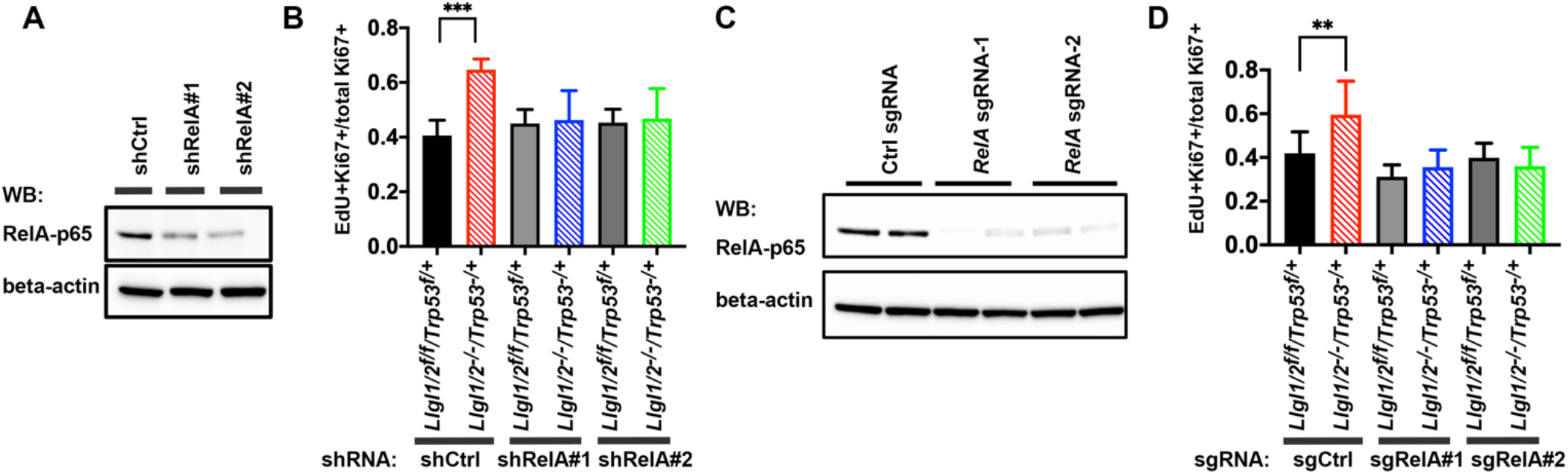
*RelA* is necessary for hyperproliferation of *Llgl1/2*^*-/-*^*/Trp53*^*+/-*^ primary keratinocytes. **A**. Knockdown efficiency of lentiviral shRelA constructs. Keratinocytes were stably transduced with shCtrl, shRelA#1 and shRelA#2 lentiviruses and analyzed by Western blotting with anti-RelA and anti-beta-actin antibodies. **B**. Quantitation of one hour pulse EdU incorporation of primary keratinocytes with indicated genotypes stably transduced with indicated shRNA lentiviruses. **C**. Knockout efficiency of *RelA* targeting CRISPR lentiviral constructs. Keratinocytes were stably transduced with sgCtrl, sgRelA#1 and sgRelA#2 lentiviruses and analyzed by Western blotting with anti-RelA and anti-beta-actin antibodies. **D**. Quantitation of one hour pulse EdU incorporation of primary keratinocytes with indicated genotypes stably transduced with indicated sgRNA lentiviruses. Graphs show mean values with standard deviation. ** - indicates P value <0.01. *** - indicates P value <0.001. P value was determined using Student’s t-test.

We hypothesized that activated aPKC is responsible for increased NF-kB signaling in *Llgl1/2*^*-/-*^ cells. Indeed, shRNA mediated knockdown of aPKCi erased the differences in activating phosphorylation of IKK-beta and RelA between *Llgl1/2*^*-/-*^*/Trp53*^*-/+*^ and control *Llgl1/2*^*fl/fl*^*/Tp53*^*fl/+*^ keratinocytes (Fig. 7A-C). Moreover, knockdown of aPKCi also erased the differences in EdU incorporation between these cells (Fig. 7D). Therefore, we conclude that *Llgl1/2*-loss mediated activation of aPKC-NF-kB signaling pathway is responsible for hyperproliferation of *Llgl1/2*^*-/-*^*/Trp53*^*-/+*^ keratinocytes.

**Fig. 7.**
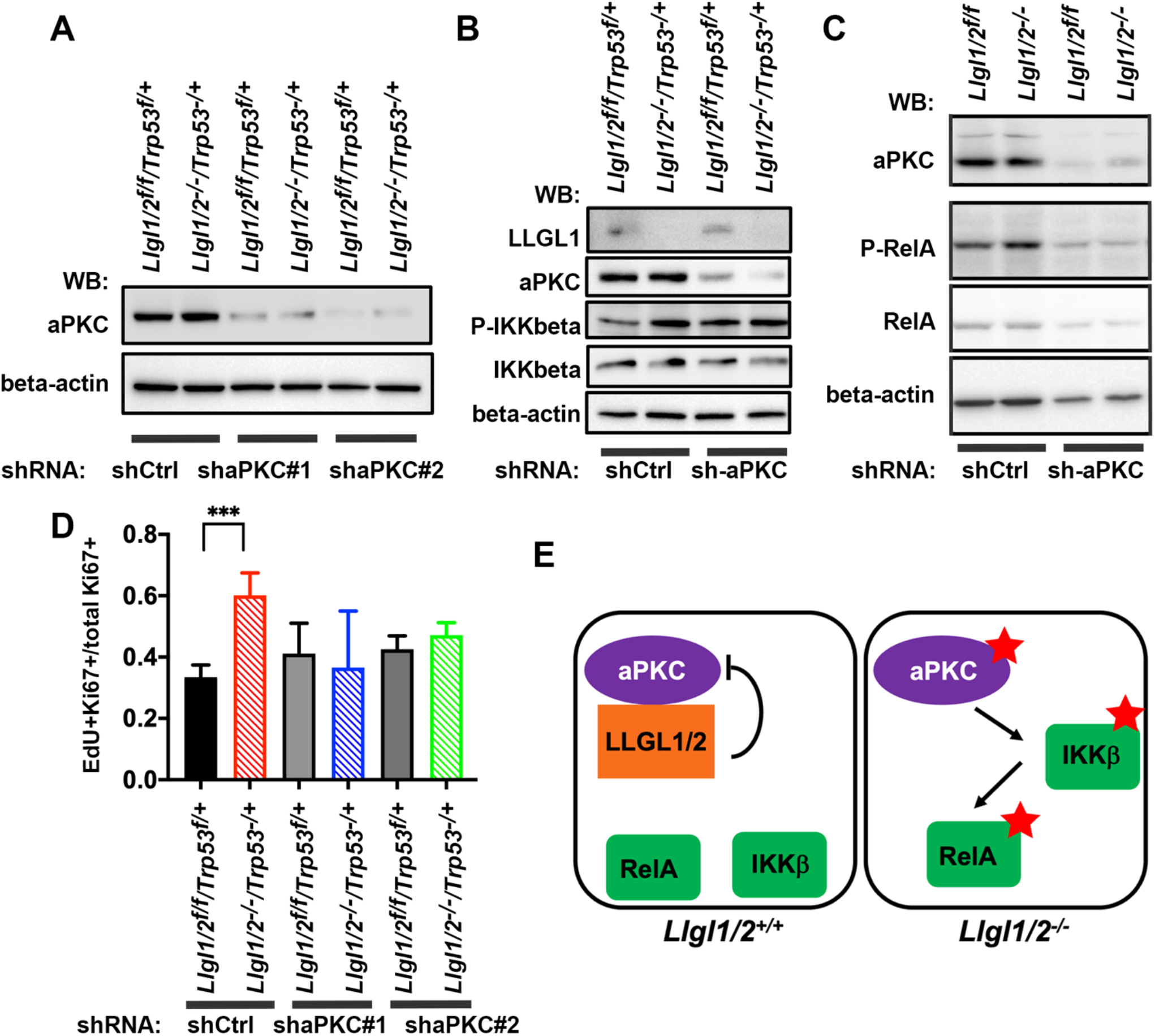
*aPKC* is necessary for hyperproliferation of *Llgl1/2*^*-/-*^*/Trp53*^*+/-*^ primary keratinocytes. **A**. Knockdown efficiency of lentiviral sh-aPKC constructs. Keratinocytes were stably transduced with shCtrl, sh-aPKC#1 and sh-aPKC#2 lentiviruses and analyzed by Western blotting with anti-aPKC and anti-beta-actin antibodies. **B**. Knockdown of aPKC erases the differences in IKK-beta activity between *Llgl1/2*^*-/-*^*/Trp53*^*+/-*^*and Llgl1/2*^*fl/fl*^*/Trp53*^*+/fl*^ controls. Keratinocytes were stably transduced with shCtrl and sh-aPKC#1 lentiviruses and analyzed by Western blotting with anti-LLGL1/2, anti-aPKC, anti-phospho-IKKbeta (P-IKKbeta), anti-total IKKbeta and anti-beta-actin antibodies. **C**. Knockdown of aPKC erases the differences in activating RelA phosphorylation between *Llgl1/2*^*-/-*^*/Trp53*^*+/-*^*and Llgl1/2*^*fl/fl*^*/Trp53*^*+/fl*^ controls. Keratinocytes were stably transduced with shCtrl and sh-aPKC#1 lentiviruses and analyzed by Western blotting with anti-aPKC, anti-phospho-RelA (P-RelA), anti-total RelA and anti-beta-actin antibodies. **D**. Quantitation of pulse EdU incorporation of primary keratinocytes with indicated genotypes stably transduced with indicated shRNA lentiviruses. Graph shows mean values with standard deviation. P value was determined using Student’s t-test. **E**. Hypothetical model of NFkB signaling activation in *Llgl1/2*^*-/-*^ keratinocytes.

## DISCUSSION

*Lgl* in Drosophila regulates asymmetric cell division of stem cells and functions as a tumor suppressor (Bilder et al. 2000). Since mammalian genomes contain two drosophila *Lgl* homologues (*Llgl1* and *Llgl2*) that have different expression patterns and different mutant phenotypes, it has been difficult to assess whether the overall function of the Lgl signaling pathway is consistent between flies and mammals (Klezovitch et al. 2004; Sripathy et al. 2011). Indeed, the published data are somewhat confusing and fragmented. Depending on the model system, *Llgl1* and *Llgl2* were strongly implicated in both tumor suppressive and tumor promoting functions (Hawkins et al. 2014; Li et al. 2017; Daynac et al. 2018; Saito et al. 2019). To begin to address the role of the entire Lgl signaling pathway in mammalian cancer, in this study we decided to generate mouse mutant with inactivation of both *Llgl1* and *Llgl2* genes. This work revealed that Lgl signaling functions as a tumor suppressor in mammalian epidermis. This finding is consistent with extensively characterized role of *Lgl* in Drosophila model system indicating the conservation of *Lgl* function between flies and mammals (Bilder 2004).

To determine the potential mechanisms of tumor-suppressive function of Lgl signaling pathway in skin epidermis we performed unbiased RNA-Seq analyses. Activation of NF-kB signaling in *Llgl1/2*^*-/-*^ epidermis and cultured primary keratinocytes was the most notable change revealed by these experiments. This was completely unexpected, because Lgl signaling has not been previously implicated in regulation of NF-kB pathway. In fact, previously published information concerning the molecular mechanisms of LLGL1/2 in regulation of cell proliferation was somewhat incoherent, with regulation of Notch signaling (Portela et al. 2015), Hippo pathway (Grzeschik et al. 2010; Parsons et al. 2014a; Greenwood et al. 2016), integrin β1 (Ma et al. 2022), MET, ErbB, and EGFR tyrosine kinase signaling (Reischauer et al. 2009; Kashyap et al. 2013; Greenwood et al. 2016), and ubiquitin-mediated degradation (Yamashita et al. 2015) proposed as potential mechanisms in different studies. Here we provide strong in vivo evidence that the NF-kB signaling is the critical mechanism of Lgl function in mammalian organism. Moreover, our experiments in primary cells indicate that this is a cell autonomous and important for regulation of cell proliferation Lgl function.

We have also determined that negative regulation of aPKC is the critical function of mammalian Lgl signaling pathway which is responsible for NF-kB regulation. Connection between Lgl and aPKC has been well established and this function is responsible for Lgl-mediated tumor suppressive function in Drosophila, although it is not known whether it also involves the NF-kB pathway downstream of aPKC (Lee et al. 2006). While we found that NF-kB signaling downstream from aPKC is critical for LLGL1/2-mediated regulation of cell proliferation in cells in culture, it is not clear if NF-kB is indeed a signaling pathway that is responsible for Lgl tumor suppressive function in vivo. Future in vivo studies will help to address this question.

While we found that activation of NF-kB was important for Lgl function in regulation of cell proliferation, it is curious that this was evident only in cells with inactivation of one allele of *Trp53*. We used *Trp53* in our studies, because *Trp53* is a relevant for skin epidermis tumor suppressor which is altered in 70% of human skin SCC tumors (Inman et al. 2018). Interestingly, Tp53 is one of the tumor suppressor genes that shows strong haploinsufficiency, where the loss of only one allele is functionally significant (Berger and Pandolfi 2011; Winiecka-Klimek et al. 2014). For example, *Trp53*^*+/-*^ thymocytes display impaired apoptosis (Clarke et al. 1993) Overall, reduction in TP53 dosage and function can impact cellular functions that play an important role in tumor suppression (Berger et al. 2011). If deletion of *Llgl1/*2 results in both increased proliferation and increased apoptotic cell death, it is possible that decreased expression of TRP53 in *Trp53*^*+/-*^ keratinocytes rescues the apoptotic part of Lgl function and thus reveals its role in stimulation of cell proliferation and tumor suppression. Future studies will have to experimentally address this potential explanation of *Llgl1/2* and *Trp53* cooperation in tumor suppression.

## MATERIALS AND METHODS

### Animal Models

All procedures involving mice and experimental protocols were approved by the IACUC of Fred Hutchinson Cancer Center (FHCC) and followed NIH guidelines for animal welfare. Mice with a conditional *Llgl1* allele (Llgl1^tm1Vv^) were previously generated in our laboratory (Jossin et al. 2017). R1 ES cells with *Llgl2* gene trapped using UPA vector (clone # 422C9, Llgl2^Gt(422C9)Cmhd^) were obtained from Toronto Centre for Phenogenomics. *Llgl2* mutant mice were generated by microinjection of ES cells into mouse embryo using conventional embryonic stem cell technology. PCR with oligos Lgl2trap2b-3′ (5′-caggcgcataaaatcagtca-3’), Lgl2trap2e (5′-gaagagtgaggtagaatctc-3′) and Lgl2trapeA (5′-cactctgctggatgacaata-3’) was used for genotyping (*Llgl2*^*+*^ wild-type allele, 0.57 kB; *Llgl2*^*fl*^ floxed allele, 1kB). Conditional floxed *Trp53* mice (Trp53^tm1Brn^) were obtained from NCI repository (stock # 01XC2).

Crossing over was used to place *Llgl1* and *Llgl2* mutant alleles on the same chromosome. For this purpose, mice with homozygous *Llgl1* allele were crossed with animals with homozygous *Llgl2* allele and resulting double heterozygous mice were crossed again to *Llgl1*^*fl/fl*^ *mice*. Progeny with crossing over event were genotyped as *Llgl1*^*fl/fl*^*/Llgl2*^*fl/+*.^ These animals were crossed to each other to obtain double homozygous *Llgl1*^*fl/fl*^*/Llgl2*^*fl/fl*^ mice. Similar approach was used to generate triple mutant *Llgl1*^*fl/fl*^*/Llgl2*^*fl/fl*^*/Trp53*^*fl/fl*^ *and Llgl1*^*fl/fl*^*/Llgl2*^*fl/fl*^*/Trp53*^*+/fl*^ mice.

For skin epidermis specific gene targeting, mutant *Llgl1/2, Llgl1/2/Trp53* and *Trp53* mice were crossed with our K14-Cre animals (Vasioukhin et al. 1999). All mice were maintained on a mixed 129S1/SvlmJ / C57BL/6J/ genetic background.

### Tissue dissection, histology and immunohistochemistry

For paraffin sections, the new born skins were fixed in 4% formaldehyde in PBS overnight, processed and embedded in paraffin. Sections (5μm thick) were stained and imaged using a Nikon TE 200 microscope. For cryosections, tissues were frozen in OCT and sectioned (7μm thick) using a Leica cryostat. For histology, sections were stained with hematoxilin & eosin. For immunofluorescent staining, skin sections were deparaffinized, rehydrated, and antigenic sites were unmasked using either Tris-EDTA (10mM Tris-HCl, pH 9.0; 1mM EDTA; 0.05% Tween-20) or citric acid-based unmasking solution (Vector Laboratories) in Pascal pressure chamber (Dako). The sections were immunostained using EnVision and ARK kits (DAKO, K400311-2 and K395411-8) according to manufacturer protocols. For immunofluorescent staining of frozen sections, slides were incubated in 4% paraformaldehyde for 15 minutes. Cells were permeabilized with 1xPBS, 0.1% Triton X-100 for 15 minutes and incubated with primary antibodies overnight at 4° C. Primary antibodies were detected using anti-mouse-conjugated Texas Red (Jackson ImmunoResearch, 115-075-075) and anti-rabbit-conjugated fluorescein isothiocyanate (FITC) (Jackson ImmunoResearch, 711-095-152) secondary antibodies. Sections were mounted using mounting medium with DAPI (Abcam, ab104139) and imaged using confocal laser scanning microscope LSM 800 (Zeiss).

### Primary mouse keratinocyte cell isolation and culture

Primary mouse keratinocytes were isolated from new-born pups as previously described (Vasioukhin et al. 2000). 1×10^6^ cells were plated on one well of 6 well-plate previously coated with 10*μ*g/ml of laminin (Sigma, #L2020) in 1xPBS for 1 hour at 37C. Keratinocytes were cultured in Keratinocyte serum-free media (KSFM) containing: Advanced DMEM/F-12 media (Thermo Fisher, 12634-010), 1x B27 Supplement (Thermo Fisher, 17504-001), 10 mM HEPES, 2mM GlutaMAX (Thermo Fisher, 35050-061), 1.25 mM N-Acetyl-L-cysteine (Sigma, A9165), 1 μM prostaglandin E2 (Tocris, 2296), 10 mM Nicotinamide, 100 μg/ml primocin, 50 ng/ml EGF (Peprotech, 315-09), 500nM A83-01 (Tocris, 2939), 500 ng/ml R-Spondin (Peprotech, 120-38) and 100 ng/ml Noggin (Peprotech, 120-10C). Media was replaced every 2 days. Cells were passaged 1:5. To passage, cells were incubated with dispase (Stem Cell Technologies, 07913) containing 10 μm Y-27632 for 5 minutes at 37C, disrupted by pipetting and transferred to 15mL conical tube. Cells were spinned down, washed once in KSFM and plated on laminin coated plates To generate paired *Llgl1/2*^-/-^, *Llgl1/2*^-/-^*/Tp53*^-/+^ *and Trp53*^-/+^ and corresponding control primary mouse cultures, keratinocytes from *Llgl1/2*^*fl/fl*^, *Llgl1/2*^*fl/fl*^ */Tp53*^*fl/+*^ *and Trp53*^*fl/+*^ mice were infected with adenoviruses carrying Cre (Ad5CMVCre-eGFP) or GFP as control (Ad5CMVeGFP), which were purchased from the University of Iowa Viral Vector Core Facility. Knockout efficiency was validated by PCR and western blot analysis.

### Cell line culture and lentivirus production

Human embryonic kidney 293T (HEK293T) cells were purchased from ATCC and cultured in DMEM media (Thermo Fisher, 11965-092) with 10% fetal bovine serum (Hyclone), sodium pyruvate (Thermo Fisher, 11360-070), non-essential amino acids (Thermo Fisher, 11140-050) and primocin (InvivoGen). Lentiviruses were produced in HEK293T cells as described (Lois et al. 2002).

### Western blot analysis and Immunofluorescent staining of cultured keratinocytes

For western blot analysis, cells or mouse newborn epidermises were lysed on ice using RIPA buffer RIPA buffer [50 mM Tris pH 8.5, 150 mM NaCl, 1%NP-40, 0.5% SDS, 1 mM EDTA, 1 mM DTT] containing a protease (Thermo Fisher, A32955) and phosphatase (Thermo Fisher, A32957) inhibitor cocktails. Protein extracts containing equal amounts of protein (50 μg) were solubilized in 1xLDS Sample Buffer (Thermo Fisher, NP0007) and separated by sodium dodecyl sulfate/ polyacrylamide gel electrophoresis (SDS-PAGE), and transferred to PVDF membranes (Millipore, IPVH00010). Membranes were incubated with primary and species-specific HRP-labeled secondary antibodies (Jackson ImmunoResearch Laboratories), that were detected using immobilon western chemiluminescent HRP substrate (Millipore, WBKLS0500).

For EdU incorporation experiment 1.0 ×10^4^ primary keratinocytes were plated in triplicates on glass cover sleeps coated with laminin. Growing in KSFM cells were incubated for 1 hour with 10mM EdU, washed with PBS, fixed for 15 mins in 3.7% formaldehyde in 1xPBS, and permeabilized with 0.5% Triton-X100 in 1xPBS. Keratinocytes were stained for EdU using Click-iT EdU kit (Invitrogen, C10340) and then blocked in Superblock (ThermoFisher Scientific, 37515) + 5% Normal Goat Serum (Jackson Immunoresearch) for 1 hour at room temperature. Cover sleeps were incubated with anti-Ki67 antibodies (1:200) overnight at 4^0^C, washed and incubated with anti-rabbit-conjugated fluorescein isothiocyanate (FITC) (Jackson Immunoresearch, 711-095-152) secondary antibodies. The coverslips were mounted on microscope slides using Mounting Medium with DAPI (Abcam, ab104139) and imaged using confocal laser scanning microscope LSM 800 (Zeiss) and quantified using image J software.

### Antibodies

For Western blotting we used the following antibodies: rabbit anti-LLGL1 (1:20,000, Sigma, HPA023569); rabbit anti-LLGL2 (1:1000, Sigma, HPA022913); rabbit anti-RelA-p65 (1:1000, Cell Signaling, #3033); rabbit anti-p-RelA-p65 (1:1000, Cell Signaling, #8242); mouse anti-aPKC (1:1000, Santa Cruz, sc-17781); rabbit anti-p-IKKα/β (1:500, Cell Signaling, #2697), rabbit anti-IKKβ (1:1000, Cell Signaling, #8943), rabbit anti-p-YAP (1:500, Cell Signaling, #4911), rabbit anti-YAP (1:500, Cell Signaling, #4912), rabbit anti-LATS1/2 (1:1000, AbCam, #ab70565), rabbit anti-p-LATS1/2 (1:500, Cell Signaling, #9159), mouse anti-p-AKT (1:1000, Cell Signaling, #4051), rabbit anti-AKT (1:1000, Cell Signaling, #9272), mouse anti-P-MAPK (1:2000, Sigma #M-8159), rabbit anti-MAPK (1:500, Sigma #M-5670), rabbit anti-TAZ, (1:1000, Sigma, #HPA007415), mouse anti-beta actin (1:5000, Sigma, #A5441).

For Immunostaining, we used the following antibodies: rabbit anti-LLGL1 (cross-react with LLGL2 (Choi et al. 2019) (1:100, Abcam, #183021); rat anti-Integrin β4 chain (1:100, BD Transduction Laboratories, #553745), rabbit anti-Ki67 (1:300, Novo Castra, #NCL-Ki67p). anti-mouse-conjugated Texas Red (Jackson Immunoresearch, 115-075-075) and anti-rabbit-conjugated fluorescein isothiocyanate (FITC) (Jackson Immunoresearch, 711-095-152).

### Plasmids

The lentivirus shRNA plasmids targeting mouse *RelA* and *Pkci* were generated by cloning annealed oligos into AgeI and EcoRI sites of pLKO.1-hygro-ctrl plasmid (Addgene #24150). Following target sequences and oligos were utilized: shRNA RelA 1 <u>TRCN0000244319</u> with target sequence GCATGCGATTCCGCTATAAAT. Forward sequence: 5’- CCGGGCATGCGATTCCGCTATAAATCTCGAGATTTATAGCGGAATCGCATGCTTTTTG-3’

Reverse sequence: 5’- AATTCAAAAAGCATGCGATTCCGCTATAAATCTCGAGATTTATAGCGGAATCGCATGC-3’ shRNA

RelA 2 <u>TRCN00000555343</u> with the target sequence GCGAATCCAGACCAACAATAA. Forward sequence: 5’- CCGGGCGAATCCAGACCAACAATAACTCGAGTTATTGTTGGTCTGGATTCGCTTTTTG-3’

Reverse sequence: 5’- AATTCAAAAAGCGAATCCAGACCAACAATAACTCGAGTTATTGTTGGTCTGGATTCGC-3’ shRNA

PKCi 1 <u>TRCN0000219728</u> with target sequence CTTCATGAGCGAGGGATAATT. Forward sequence: 5’- CCGGCTTCATGAGCGAGGGATAATTCTCGAGAATTATCCCTCGCTCATGAAGTTTTTG-3’

Reverse sequence: 5’- AATTCAAAAACTTCATGAGCGAGGGATAATTCTCGAGAATTATCCCTCGCTCATGAAG-3’ shRNA

PKCi 2 <u>TRCN0000022755</u> with target sequence CGAGGGATAATTTATAGAGAT. Forward sequence: 5’- CCGGCGAGGGATAATTTATAGAGATCTCGAGATCTCTATAAATTATCCCTCGTTTTTG-3’ Reverse sequence: 5’-

AATTCAAAAACGAGGGATAATTTATAGAGATCTCGAGATCTCTATAAATTATCCCTCG-3’. Non-targeting shRNA plasmid pKLO.1-puro (Sigma, SHC202) and empty pKLO.1-hygro (Addgene, 24150) were used as negative controls.

Mouse *RelA* sgRNA target sequences were designed using the Broad Institute sgRNA designer tool (portals.broadinstitute.org/gpp/public/analysis-tools/sgrna-design). Plasmids with sgRNA target sequences cloned into pLenti-U6-sgRNA-SFFV-Cas9-2A-Puro vector were obtained from ABM Inc. Following target sequences were utilized: *RelA* sgRNA1: 5’-GATTCCGCTATAAATGCGAG-3’ *RelA* sgRNA2: 5’-GGTCTGGATTCGCTGGCTAA-3’, *RelA* sgRNA3: 5’-GTTCCTATAGAGGAGCAGCG-3’. Scrambled sgRNA vector was used as negative control (ABM Inc., K010)

### RNA extraction and RNA-Seq analysis

Total RNA was extracted with QIAzol (QIAGEN) followed by RNase-free DNAase treatment (QIAGEN) and purification using RNeasy kit (QIAGEN). RNA concentration, purity, and integrity was assessed by NanoDrop (Thermo Fisher Scientific Inc) and Agilent TapeStation. RNA-seq libraries were constructed from 1 ug total RNA using the Illumina TruSeq Stranded mRNA LT Sample Prep Kit according to the manufacturer’s protocol. Barcoded libraries were pooled and sequenced on a NovaSeq S1 100 flowcell generating 50 bp paired end reads. Sequencing reads were mapped to the mm10 mouse genomes using STAR.v2.7.3a1. Gene level abundance was quantitated using GenomicAlignments (Lawrence et al. 2013) and analyzed using limma(Ritchie et al. 2015), filtered for a minimum expression level using the filterByExpr function with default parameters prior to testing, and using the Benjamin-Hochberg false discovery rate (FDR) adjustment. Genome-wide gene expression results were ranked by their limma t-statistics and used to conduct Gene Set Enrichment Analysis (GSEA) to determine patterns of pathway activity utilizing the curated pathways from within the MSigDBv7.4(Subramanian et al. 2005).

### Statistical Analyses

Statistical significance was determined by Student’s, Fisher’s exact, Mann-Whitney, or Chi-square tests. p value is indicated by asterisks in the figures: *p < 0.05; **p < 0.01; ***p < 0.001. Differences at p = 0.05 and lower were considered statistically significant. In RNA-seq analyses, the differences with FDR < 0.05 were considered statistically significant.

## AUTHOR CONTRIBUTIONS

Conceptualization, V.B., and V.V.; Methodology, V.B., D.R., O.K. and V.V.; Investigation, V.B., D.R. and O.K.; Writing – Original Draft, V.V.; Writing – Review & Editing, all authors; Funding Acquisition, V.B. and V.V.; Supervision, V.V.

## ACKNOWLEDGEMENTS

We thank the Comparative Medicine and Genomics Core Facility of FHCC for help with ES cell microinjection, animal care and next generation sequencing. We thank Luan T. Phan, Elizabeth Zimmerman, Bridget Kreger, Olga Chen, Alice S. Woo, Laura Sherer, Parul Katosh and Smitha Sripathy for help with this study. This research was supported in part by NCI grants R01 CA234050 and T32 CA009657, ACS grant 50000784, FHCC discretionary funds and NIH/NCI Cancer Center Support Grant P30 CA015704.

## SUPPLEMENTARY FIGURES

**SFig.1.**
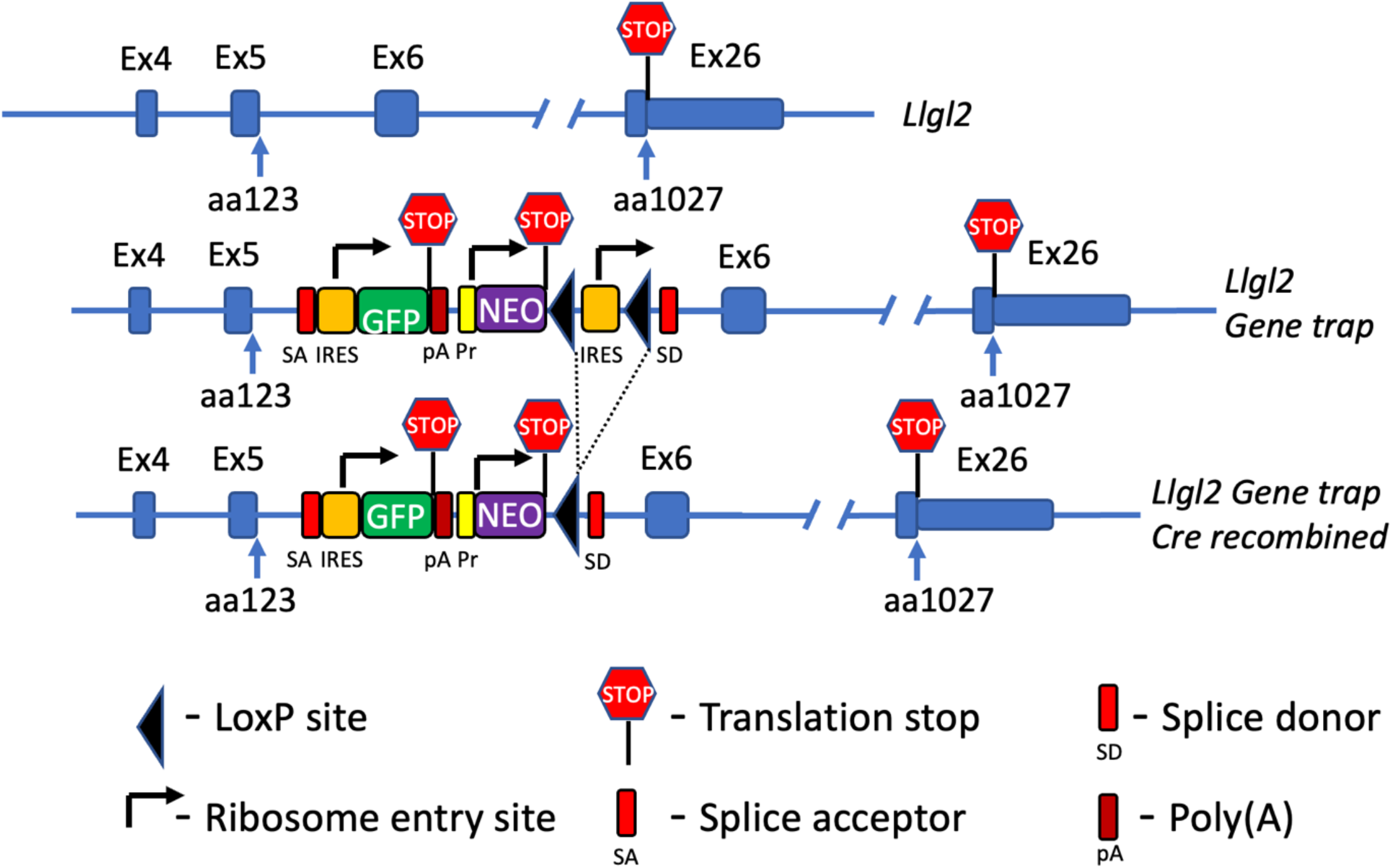
Generation of *Llgl2* mutant allele.

**SFig. 2.**
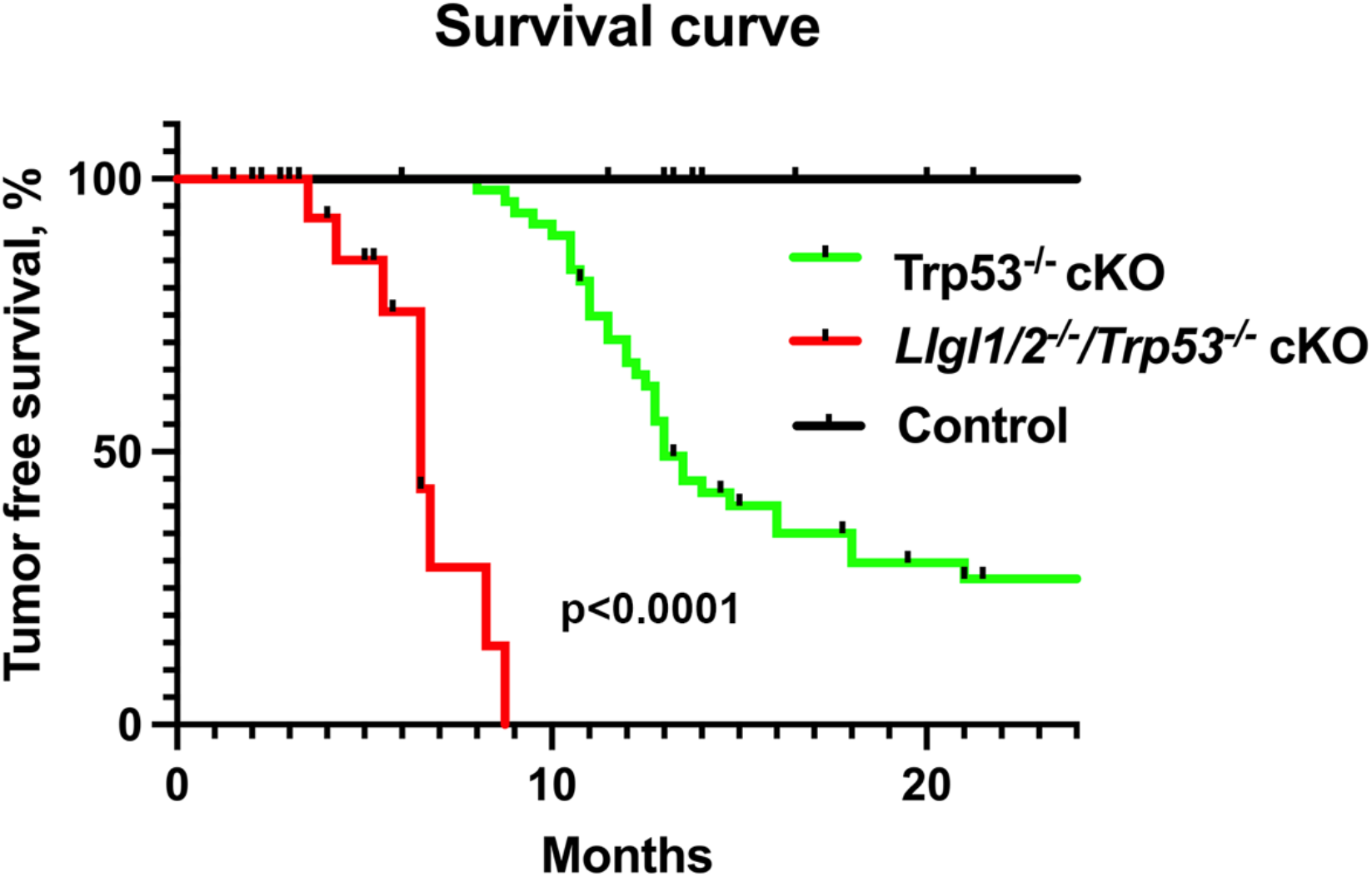
Loss of *Llgl1/2* promotes skin SCC development in mice with homozygous deletion of *Trp53*. Survival curve of Control (K14-Cre), *Trp53* cKO (*K14-Cre/Trp53*^*fl/fl*^) and *Llgl1/2*^*-/-*^*/Trp53*^*-/-*^ cKO (*K14Cre/Llgl1/2*^*fl/fl*^*/ Trp53*^*fl/fl*^) mice. Log-Rank test.

